# Predicting molecular subtypes of breast cancer using pathological images by deep convolutional neural network from public dataset

**DOI:** 10.1101/2020.02.12.946723

**Authors:** Nam Nhut Phan, Chi-Cheng Huang, Eric Y Chuang

**Affiliations:** Bioinformatics Program, Taiwan International Graduate Program, Institute of Information Science, Academia Sinica, Taipei; Graduate Institute of Biomedical Electronics and Bioinformatics, National Taiwan University, Taipei; Comprehehsive Breast Health Center, Taipei Veterans General Hospital, Taipei City, Taiwan; Biomedical Technology and Device Research Laboratories, Industrial Technology Research Institute, Hsinchu

## Abstract

Breast cancer is a heterogeneously complex disease. A number of molecular subtypes with distinct biological features lead to different treatment responses and clinical outcomes. Traditionally, breast cancer is classified into subtypes based on gene expression profiles; these subtypes include luminal A, luminal B, basal like, HER2-enriched, and normal-like breast cancer. This molecular taxonomy, however, could only be appraised through transcriptome analyses. Our study applies deep convolutional neural networks and transfer learning from three pre-trained models, namely ResNet50, InceptionV3 and VGG16, for classifying molecular subtypes of breast cancer using TCGA-BRCA dataset. We used 20 whole slide pathological images for each breast cancer subtype. The results showed that our scale training reached about 78% of accuracy for validation. This outcomes suggested that classification of molecular subtypes of breast cancer by pathological images are feasible and could provide reliable results

## 1. Introduction

Breast cancer is the most common female malignancy in Taiwan and treatment outcomes have been improved enormously in the past decade, mainly with the wide applications of screening mammography (early detection at pre-clinical stage) and advancement in adjuvant/neoadjuvant therapy. Nowadays the selections of adjuvant therapy are determined from certain clinical predictive (also prognostic) factors such as estrogen receptor (ER) and human epidermal growth factor receptor type II (HER2) status. These factors not only determine which adjuvant therapy should be executed but also predict therapeutic responsiveness.

These conventional pathological factors, however, do not fully explain the prognostic heterogeneity observed within each clinical stratum [1]. For example, one-fourth of HER2 over-expressed breast tumors eventually develop resistance to trastuzumab while hormone manipulation therapy alone is not adequate for a subset of aggressive ER positive (luminal B) breast cancer. On the other hands, microarrays have subdivided breast cancers into molecular subtypes in terms of transcriptome profiles, such as the ‘intrinsic subtype’ proposed by the Stanford/University of North Carolina group. Initially Perou et al. identified 476 intrinsic genes from 65 breast cancers and healthy individuals; four subclasses: basal-like, Erb-B2+, normal breast-like, and luminal epithelial/ER+ were revealed through clustering analysis [1,2]. The luminal subtype was further divided into luminal A and B, with the latter experienced worse survival [3]. Intrinsic genes were selected genes with the highest variations among different breast cancer patients but with the least variations within the same individual. Different generations of intrinsic signatures followed, and the latest prediction analysis of microarray 50 gene set (PAM50) was advised to provide prognostic and predictive values independent of traditional prognostic factors [4,5]. The rationale underpinning the merits of molecular biomarkers rather than conventional pathological factors may result from that transcriptional aberrations essential for breast cancer pathogenesis are investigated, while clinical predictors are merely manifesting phenotypes.

During the 2011 St. Gallen experts’ panel, immunohistochemical (IHC) surrogates for breast cancer molecular subtypes using tumor grade instead of Ki-67 had demonstrated that gene expression microarrays-defined breast cancer molecular subtypes could be approximated by conventional IHC assays [2]; hormone receptor positive breast cancers are categorized into luminal A (HER2- and Ki-67<14% or nuclear grade I/II), luminal B1 (HER2- and Ki-67>14% or nuclear grade III), and luminal B2 (HER2+). Luminal B breast cancer tends to display a more aggressive and compromised clinical outcomes, and this subtype is presented as ER+ tumors with low progesterone receptor expression, high proliferation, high grade, and less responsive to hormone manipulation therapy [3].

Artificial intelligence (AI) plays a crucial role in biomedical image analysis and cancer research. Certain breast carcinomas behave aggressively resulting in increased patient morbidity and poor patient prognosis. There are distinguishable cytological features of such tumors including aggressive variants of hereditary breast carcinoma, poorly differentiated metaplastic carcinoma, and triple negative breast cancer. *BRCA1-associated* breast cancers are commonly poorly differentiated, have “medullary features” (a syncytial growth pattern with pushing margins and a lymphocytic response), and are biologically very similar to basal-like subtype defined by gene expression profiling. *BRCA2*-associated breast cancers also tend to be relatively poorly differentiated but are more often ER-positive than *BRCA1* mutant counterparts. With the utility of digitalized, whole-slide images, it is possible to develop an algorithm by artificial intelligence. A machine-learning algorithm may be applied to histopathological images of breast cancer specimens to see if it could pick out distinguishing patterns. Therefore, it is an urgent necessity in applying computer aided enhancement of breast pathological imaging diagnosis which could augment the performance of diagnostic accuracy, reducing both false positive results and negativities. Consequently, it would be of great help to scientific communities and clinical practitioners if molecular gene expression-defined molecular subtypes could be identified by pathology pattern recognition. In the present study, we aim to classify the molecular subtypes of breast cancer via pathogical images by deep convolutional neural network and transfer learning with pre-traine model such as ResNet50, InceptionV3, and VGG16.

## 2. Materials and methods

Two training strategies for specific subtype classification were established. The former approach used patches generated from whole slides image as training data. The latter would use an althernative approach which mounts the whole slide image for training (Figure 1). The second approach would only require gene expression assay results for the final molecular subtyping, thus it may help pathologists spend less time in annotation.

**Figure 1.**
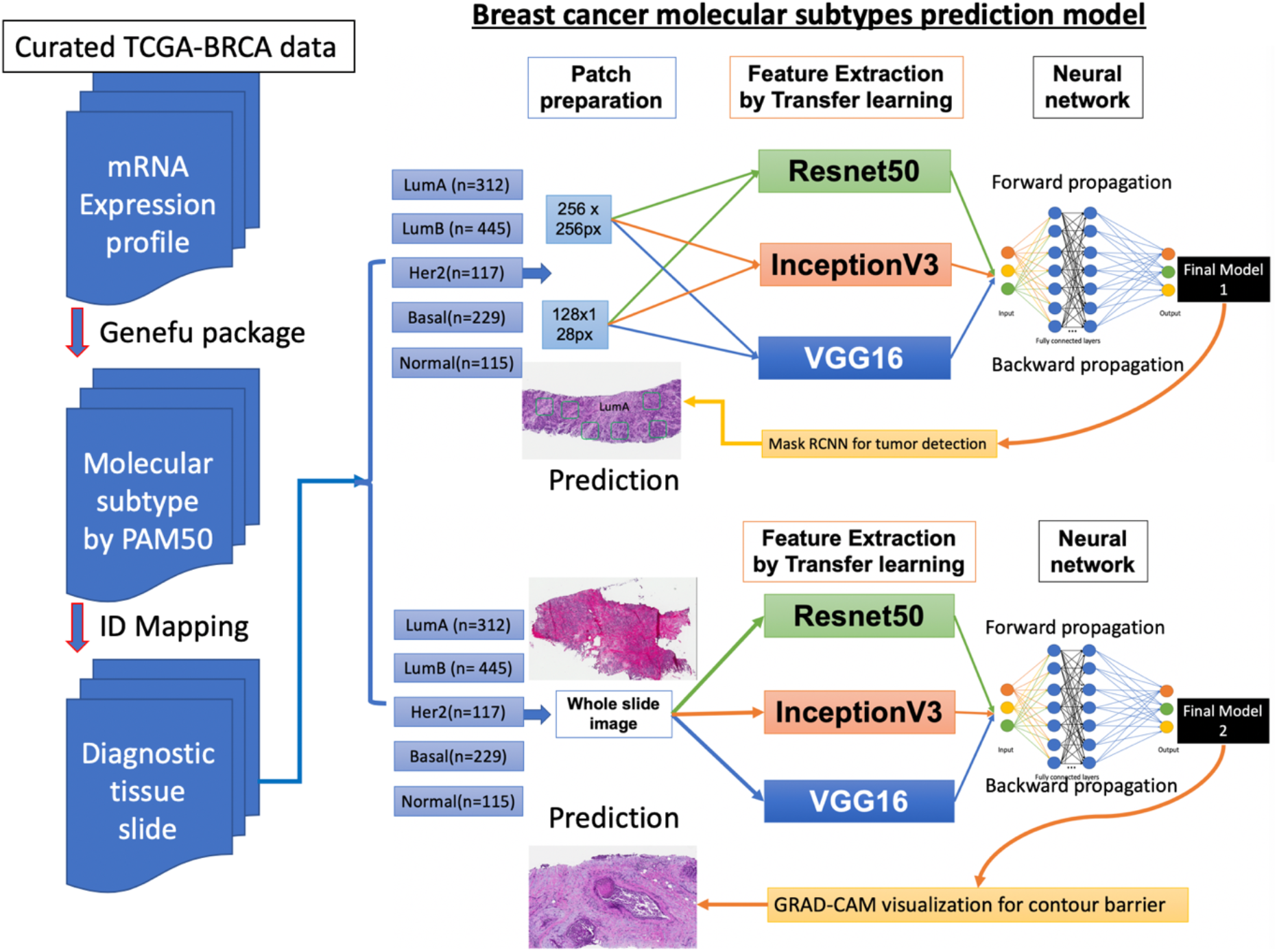
Design scheme for molecular subytpes training. The first approach used transfer learning and Mask Recurent Convolutional Neural Network (RCNN) design for tumor detection with patches generated from whole slide image as samples.

Regarding molecular subtypes, 1218 whole slide images from TCGA-BRCA dataset were used to classify patients based on their mRNA expression profiles using genefu package in R software. The mRNA expression data of TCGA-BRCA was obtained from UCSC Xena database (https://xenabrowser.net/datapages/). We designed 2 strategies for the training based on different types of samples input including whole slide image based and patches-based. The former used whole slide image from diagnostic slides whereas the latter used patching method to generate patches at 256×256 and 128×128 pixel size. These samples were the input for transfer learning process and finally undergone artificial neural network for classification and output the prediction result. This model output is the subtypes of pathological images and their tumor locations.

### 2.1 Patch generation

A total of 1218 whole slides images from distinct molecular subtypes of breast cancer underwent patch generation with py-wsi (https://github.com/ysbecca/py-wsi). Py-wsi is a python-based tool allowing us to query the whole slide resolution at different level, the level tile and level dimension. In our study, we use py-wsi package to generate patches size of 256×256 pixel and 128×128 pixel at level 17 which is the highest resolution level.

### 2.2 Whole slide image preparation

The whole slide tissue were scanned with Aperio scanncer (Leica Biosystems, IL) to generated the digital file with .svs format. The .svs files were mounted at the highest resolution and used as training samples.

### 2.3 General design

The general design of our deep convolution neural network included two stages. The first stage applied transfer learning techniques which take advantages of state of the art pre-trained models such as ResNet50 [4], InceptionV3 [5], and VGG16 [6] for potential useful features extraction []. The second stage was training with our designed hidden layers containing four fully connected layers with 512 neurons in each layer which connected with the final output layer using the softmax activation function. After obtaining highly robust model, we can used these model for classifying different subtypes of breast cancer. The mask Recurent Convolutional Neural Network (RCNN) [7] and Gradient-weighted Class Activation Mapping (Grad-CAM) [8] output were used for vizualization for model 1 and model 2, respectively.

Experiment design using transfer learning for features extractions with three models namely ResNet50, InceptionV3, and VGG16 models followed by pre-designed hidden layers for classification training. Transfer learning and Gradient-weighted Class Activation Mapping (Grad-CAM) visualization for contour barrier detection with whole slide image as training sample was the second approach.

### 2.4 Transfer learninng

Transfer learning with ResNet50 and VGG16 was carried out. The top layers of these models were not included in the feature extraction process, only convolutional layers were used. The input images were reshaped as default setting at 224×224 pixel for ResNet50 and VGG16.

The feature files were exported with different dimension depending on model output. The output dimensions of each model were displayed in Table 1.

**Table 1.** Features dimensions produced by VGG16, ResNet50 and InceptionV3 with patch-size of 256×256 pixel and 128×128 pixel.

| Patches size | VGG16 | ResNet50 | InceptionV3 |
| --- | --- | --- | --- |
| 256x 256 pixel | 7*7*512 | 7*7*2048 | 6*6*2048 |
| 128x128 pixel | 4*4*512 | 4*4*2048 | 2*2*2048 |

### 2.5 Training procedures

Data were divided into 80% for training (training and validation set) and 20% for testing. The testing data was only used when the training was completed to prevent information leaking to the network.

The feature map files from Resnet50, InceptionV3 and VGG16 were used as input to artificial neural network with 4 hidden layers, each contains 512 neurons. The final connected layer linked to the output layer with 5 neurons, where each neuron represented one molecular subtype. We used stochastic gradient descent (SGD) as the optimizer with learning rate at 10e^-5^ together with a decay rate 10e^-5^/10 for 10 epochs with the batch size set to 128. Kernel was used with rectified activation unit (<u>ReLu</u>) function with kernel initnitializer was “he_uniform”. Kernel regularizer was used with L2 at 0.0001. To prevent model overfitting, we applied dropout techniques at 0.2. the final dense layer was used with softmax function for final class decision. We used categorical cross entropy to calculare the loss of our model prediction and monitor the training process by categorial accuracy metric.

Owing to the massive amount of samples, we used fit generator to train our models with steps per epoch equal total number of training samples devided by batch size. The validation step was calulated by the total number of validation samples divided by batch size. We also used model checkpoints to obtain the highest performance model with callbacks. The total number of trainable parameters was 17,574,405.

At the end of training, we used predict_generator to test the model perfomance with test data. The classification reported from scikit learn[9] was used to calculate and display the final classifications of each breast cancer molecular subtype with precision, recall, and F1-score. All training was done with Tensorflow version 2.0 (Google Inc., Mountain View, CA). The hardware system contained 2 GPU GeForce GTX 1080 Ti.

## 3. Results

We obtained mRNA expression values from public domain database for molecular subtyping classifications including luminal A (n = 312), luminal B (n = 445), HER2-enriched (n = 117), basal-like (n = 229), and normal breast-like (n =115) breast cancers. These subtypes were matched to corresponding pathological diagnostic images using case ID. At the pilot study, we used 20 samples from each subtype to generate patches at 256×256 and 128×128 pixel size. The patches generated from 128×128pixel size were 894.609 patches for basal-like, 1,326,418 patches for HER2-enriched, 1,134,517 patches for luminal A, 907,278 patches for luminal B, and 1,080,685 patches for normal breast-like subtype (Table 2). These patches were fetched into the ResNet50 for feature extraction and trained with artificial neural network.

**Table 2.** Training, validation, and testing data at patch size 128×128 pixel for 5 molecular subtypes.

| Subtypes | Training | Validation | Testing |
| --- | --- | --- | --- |
| Basal-like | 715688 | 134191 | 44730 |
| Her2 | 1061136 | 198962 | 66320 |
| Luminal A | 907615 | 170177 | 56725 |
| Luminal B | 725824 | 136091 | 45363 |
| Normal breast-like | 864549 | 162102 | 54034 |

Performance of the pilot study was beyond 78% for validation accuracy with Resnet50 as the feature extractor. The results of VGG16 and InceptionV3 were still under working. We were currently working on the whole TCGA-BRCA dataset to boost the model performance.

## 4. Discussion

Different breast cancer molecular subtypes have discrepant prognoses over time. Understanding different recurrence patterns can improve breast cancer care through surveillance guidelines and result in the most optimistic treatment [10]. According to Metzger-Filho et al., basal-like and HER2+ cohorts had higher risk of recurrence in the first 4 years after diagnosis. On the other hand, luminal B had a continuously higher hazard of recurrence over 10-year follow-up compared with luminal A breast cancer. Another study by Ribelles et al. reported that luminal A displayed a slow risk increase, reaching its maximum after three years and then remained steady. Luminal B presented most of its relapses during the first five years [11]. Compared with HER2+ and triple negative cancers, ER+ breast cancers experienced more mid-to long-term relapses, and acquired ESR1 mutations, resulting in ligand-independent and constitutive activation of ER, are believed to play a major role for late recurrence and hormone manipulation therapy resistance [12].

Deciphering breast cancer molecular subtypes by deep learning approach could provide convenient and efficient way for diagnosis and treatment of breast cancer patients. It could reduce the spending budget of transcriptional profiling and subtyping discrepancy between IHC assays and mRNA expressions. In term of academic development, this is a novel approach which can lay the foundation for later research on breast cancer taxonomy and precision medicine.

We are approaching an era for artificial intelligence and machine learning-aided diagnosis and treatment. We believe that breast pathology imaging may be one of the frontiers. Artificial intelligence has the potential to transform genomics, pathology, and breast oncology to the next level, and current deep learning systems are going to match human performance in reading pathological morphological features and reducing inter-observer variability.

In current study, the first approach with ResNet50 as the feature extractor archived an overall accuracy of 78% in the testing data whereas the model with InceptionV3 had 45% accuracy. The second approach results using whole slide image will be updated in the next revision.

In previous literatures, many models have been developed to predict or classify a wide range of diagnostic or therapeutic targets of breast cancer. Cruz-Roa et al. designed a convolutional neural network to detect the location of invasive tumor in whole pathological images. In their study, they used 400 samples for training and validated the model with 200 samples from TCGA database. The model archived 75.86%, 71.62 %, and 96.77% of Dice coefficient, positive predictive, and negative predictive value relative to manual annotation of invasive ductal carcinoma [13]. In another study, Zheng et al. applied the K-means algorithm discriminating benign and malignant lesions. The feature was extracted and trained with the support vector machine (SVM). This model reached 97% of accuracy with 10-fold cross validation on Wisconsin Diagnostic Breast Cancer (WDBC) test dataset [14]. Google’s Inception model had been used for identifying cancer subtypes with extensive tumor heterogeneity with accuracy rates of 100%, 92%, 95%, and 69% for various cancer tissues, subtypes, biomarkers, and scores respectively [15]. Another study by Alakwaa et al. trained a cohort of 548 patients including breast cancers with feed-forward networks and a deep learning (DL) framework. They found that DL method had archived area under the curve (AUC) of 0.93 in classifying ER+ and ER-patients [16].

## Conclusions

In summary, our present study provided a prqctical pipeline to use pathological images by deep convolutional neural network for breast cancer molecular subtypes with high accuracy. The trained model could be used to localize the tumor location in the whole slide image. In our experiment, the model with pre-trained model namely ResNet50 had the highest accuracy.

## Acknowledgement

We thank to…

## Authors’ Contributions

### Funding

We thank to … for the funding of this project with number …

### Conflict of interst

Authors declare no conflict of interest in this study.

